# ICAM-5: A Novel Marker for Neuronally Derived Extracellular Vesicles

**DOI:** 10.1101/2025.11.07.686794

**Authors:** Piper Deleon, Reagan Hattenberger, Barbara Knollmann-Ritschel, Manish Bhomia

## Abstract

Current methods for isolating neuronal-derived extracellular vesicles (NDEVs) from human biofluids lack specificity. In addition, some of the reported markers for NDEV isolation are present as soluble proteins instead of extracellular vesicle (EV) associated proteins. To address the research gap, this study aimed to identify a novel marker for NDEV isolation that is NDEV associate and highly specific to the central nervous system. To achieve this, human cortical neurons were used for isolation of EVs *invitro*. Mass spectroscopy was performed to screen for potential EV surface markers. This analysis yielded 63 brain specific proteins among which ICAM-5 was selected for further validations in human serum and cerebrospinal fluid (CSF) samples. Our analysis shows that ICAM-5 is present on the surface of NDEVs, colocalizing with standard extracellular vesicle markers like CD-63 and CD-9. We further confirm that NDEVs isolated using ICAM-5 contain neuronal proteins (tau, neuronal specific enolase) but not glial markers. Using ICAM-5 as a NDEV marker, EVs were eluted from human serum samples of with traumatic brain injury. Our results show enhanced levels of ubiquitin C-terminal hydrolase L1 (UCH-L1), a marker of neurological injury, in serum samples from patients with TBI when compared to currently known markers of NDEV. Our findings demonstrate that ICAM-5 is specific EV associated marker for isolating NDEVs from human biofluids that can potentially improve the enrichment of NDEVs from biofluids thereby improving diagnosis and monitoring of neurological conditions.

## Introduction

Extracellular vesicles (EVs) are small vesicles that are 30 nm to 1 µm in size and are released by a variety of cell types, including cells of the central nervous system (CNS) [1]. EVs carry a diverse array of biomolecules including proteins, DNA, RNA, lipids, metabolites, and cell-surface proteins reflecting their cells of origin [1]. This unique characteristic makes EVs highly relevant as novel diagnostic tools, particularly for neurological diseases. The ability of EVs to cross the blood brain barrier (BBB) makes them ideal candidates for the discovery of biomarkers of neurodegenerative disorders [2]. Neuronal-derived EVs (NDEVs) have emerged as a promising tool for biomarker research for neurodegenerative disorders such as Alzheimer’s disease (AD), Parkinson’s disease (PD), Huntington’s disease (HD) along with CNS injuries including traumatic brain injury (TBI) and stroke. NDEVs improve biomarker specificity and sensitivity for detecting disease-related molecular changes [4]. Therefore, purifying NDEVs from biofluids such as blood, urine, and saliva is a key research challenge in detection and discovery of biomarkers for neurological disorders. Biomarker analysis for neurological disorders from peripheral blood has few limitations including a) the biomarker’s cellular origin cannot be conclusively determined, and b) low concentration of biomarkers originating in the CNS presents detection challenges. NDEVs are specific and their detection will reflect biomarkers of neuronal origin. NDEVs can be concentrated from a larger volume of plasma, which can overcome the detection limit of very low concentration of biomarkers[3]. Therefore, NDEVs can address key challenges in biomarker identification for neurological disorders.

To enrich NDEVs from biofluids, two neuronal cell surface markers have been primarily reported in literature including L1 cell-adhesion molecule (L1CAM) and Alpha-3 subunit of the Na+/K+-ATPase (ATP1A3) [3, 4]. Of these, L1CAM has been the primary marker investigated for NDEVs biomarker identification, including in our lab [7]. L1CAM is expressed in neurons, but its specificity for CNS has been questioned since it is reported to be expressed in several non-neuronal tissues [5]. In addition, reports have identified that a significant proportion of L1CAM in blood is present as a soluble protein which can lead to loss of sensitivity as well as specificity for NDEV enrichment [6, 7]. More recently, ATP1A3 has been identified as a novel surface marker for NDEVs. In neurons, ATP1A3 helps maintain the electrochemical gradients of sodium (Na+) and potassium (K+) ions across the plasma membrane. While ATP1A3 is abundantly expressed in neurons, much like L1CAM, its presence in other tissues, such as the heart and kidneys, reduces the specificity for NDEVs enrichment [8]. In addition to L1CAM and ATP1A3, Neurexin 3 (NRXN3) has also been reported as a potential marker for NDEV enrichment from biofluids [9]. Although enriched in the CNS, NRXN3 is also expressed outside of the CNS and reported to be present in the lung, pancreas, heart, placenta, liver, and kidney [10]. Therefore, there is no highly specific EV surface marker for the isolation of NDEVs from human biofluids. To address this research gap, the present study sought to identify a new marker for the enrichment of NDEVs from human biofluids. Further, comparison this novel biomarker, ICAM5, with existing NSEV markers for specificity to CNS and ability to quantitate CNS specific biomarkers is explored.

## Results

### 1. Identification of novel neuronal EV surface markers using label-free mass spectroscopy

Mass spectroscopy (MS) was used to discover new surface proteins on EVs from cultured human primary neuronal cells (PNC) (**Figure 1a**). Primary human neurons were used for this study. To assess the purity of the cells, immunofluorescence assay was performed for neuronal specific enolase (NSE), a marker for neurons, and glial fibrillary acidic protein (GFAP), a marker for glial cells. Results showed that all the cells were positive for NSE with no detectable GFAP in these cells indicating purity of the cells (**Figure 1b**).

**Figure 1.**
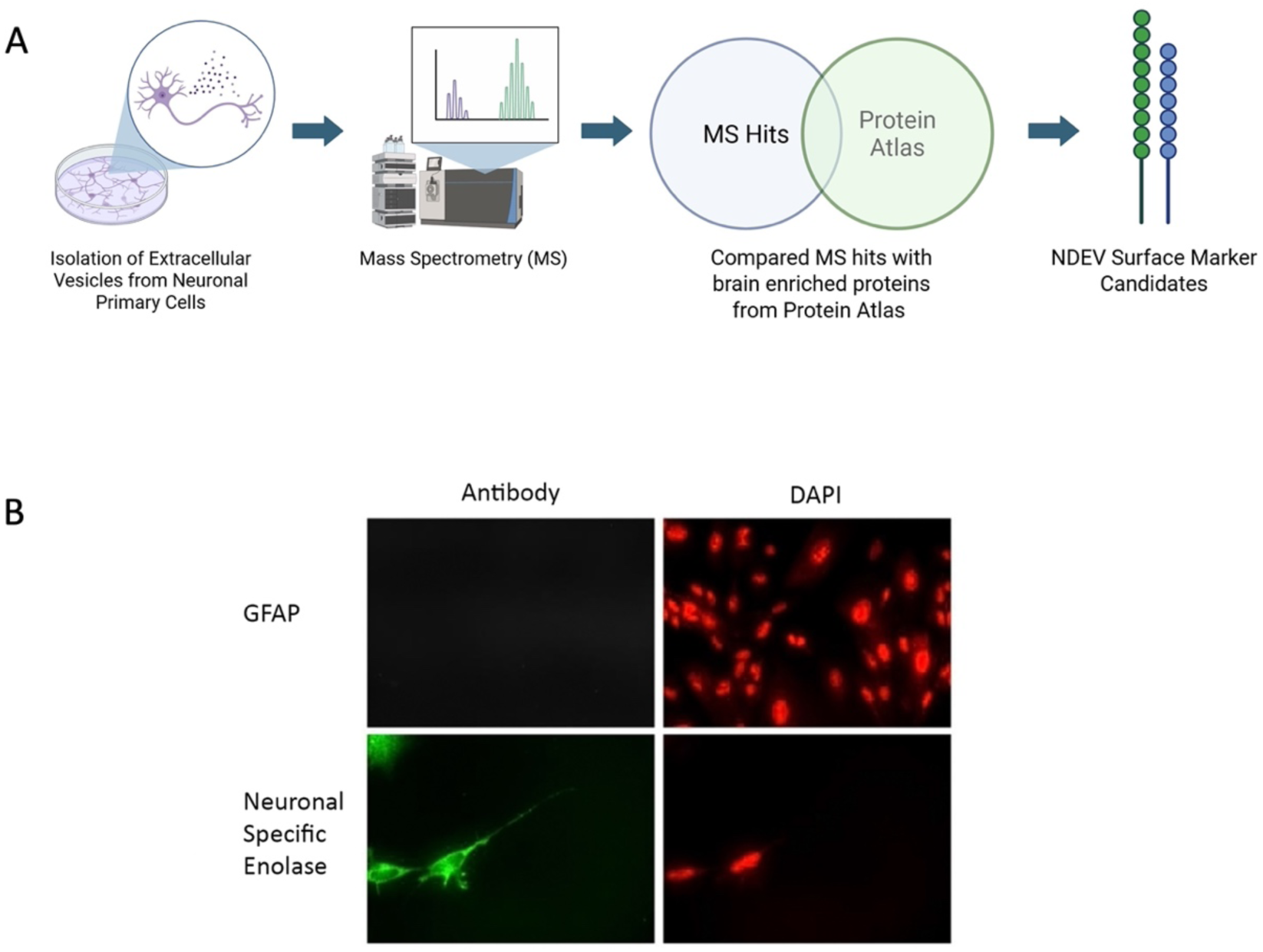
**A)** Representative workflow for discovery of novel EV markers using primary neuronal cell culture couple with mass spectroscopy (created with biorender). B) Immunofluorescence for NSE and GFAP in human neuronal primary cells. The cells showed positive staining for NSE and GFAP staining was not detected suggesting that the majority of the cells were of neuronal origin.

For MS experiments, EVs were isolated from neuronal cell culture supernatant. Prior to EV isolation, PNC culture media was replaced with serum-free 12 hr prior to EV isolation to prevent contamination from EVs present in the serum. MS analysis was performed with these EVs that yielded more than 3,000 protein hits (**Supplementary table 1**). Among these, several tetraspanins protein markers of EVs such as CD-9, CD-81 and CD-63 were identified. MS data also confirmed presence of L1CAM, NCAM1, and ATP1A3 in the overall hits; however, these proteins are not entirely specific to CNS and are expressed in other tissues. A two-step filtering criterion was used to identify proteins that are EV associate and specific to human CNS (**Figure 1a**). First, we compared the proteins identified in MS analysis the list of proteins in human protein atlas [11] under the heading “elevated proteins in the human brain”. This analysis identified 65 proteins that were present in the MS dataset (**Supplementary Table 2**). Subsequently, the 65 proteins were individually analyzed for their neuronal specificity and presence as a membrane protein that revealed one single transmembrane protein, ICAM-5.

### 2. Presence of ICAM-5 in EVs from human serum samples of mild traumatic brain injury (mTBI)

Next, the presence of ICAM-5 in EVs was confirmed in human serum samples. Total EVs isolated from pooled serum samples (N=3) from patients with a mild TBI were used. Three fractions of EVs (800 µl each) were collected and fractions two and three were pooled for these assays as they predominantly consist of EVs (**Figure 2a**). Exocheck neuro array (System Biosciences) confirmed the presence of EV specific markers including tetraspanins (CD-63, CD-9 and TSG-101) and neuronal markers (Tau, NSE and L1CAM) (**Figure 2b**). The presence of ICAM-5 was verified in these samples on a western blot with a monoclonal antibody for ICAM_5 that detected a specific band at the predicted molecular weight of approximately 130 kDa for ICAM-5 in the EVs (**Figure 2b**). Additionally, a few smaller sized bands were observed, which may represent cleaved protein. Thus, we confirmed presence of ICAM-5, along with other previously implicated NDEV markers, in EVs derived from human serum.

**Figure 2.**
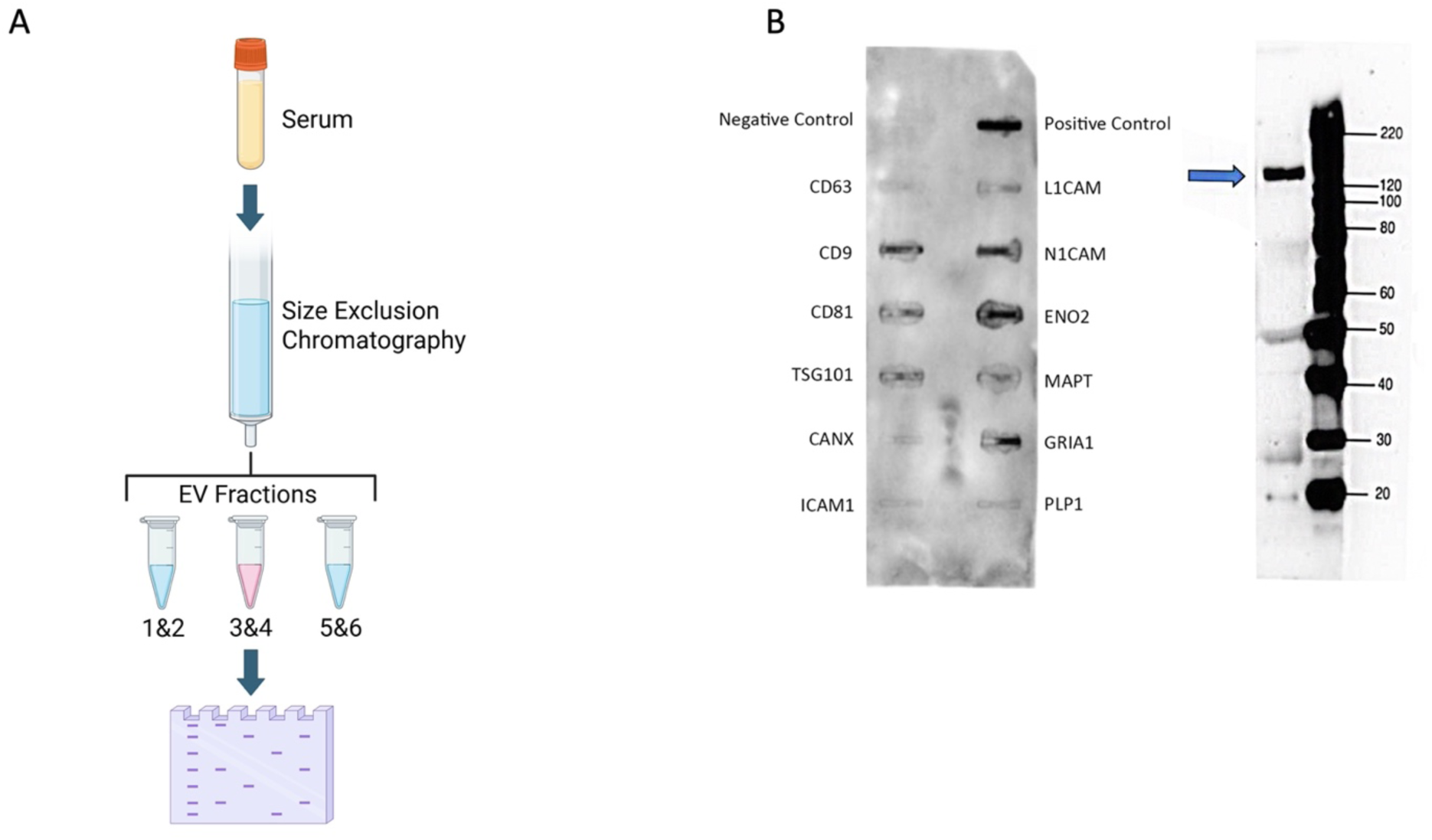
A) Schematic for EV isolation and western blot analysis (created with biorender). B) Exo-Check Exosome Antibody Neuro-Array (system Biosciences) for total EV sample isolated from pooled mTBI serum using size exclusion chromatography. B) Western blot with ICAM-5 antibody in total EVs from pooled mTBI samples.

### 3. ICAM-5 in human biofluids is associated with extracellular vehicles and is not a soluble protein

After confirming the presence of ICAM-5 in human serum, the subsequent objective was to determine if ICAM-5 primarily exists as an EV associated protein or as a soluble protein in serum. Using SEC, several EV fractions from control human serum samples were collected. If ICAM-5 was an EV-associated protein, then it would elute with EV markers such as tetraspanins CD-63 and CD-9, otherwise it will elute in the soluble protein fractions. Twenty fractions were collected from control human serum. Nanoparticles tracking analysis (NTA) confirmed the presence of EVs from fractions 7-16. This was further validated by electron microscopy (EM) where most of the EVs were only observed in fractions 7-15 (**Figure 3 a, b**). Subsequently, Western blots were performed on the serum EV fractions 3-20 and probed for CD-63, CD-9, ICAM-5, ATP1A3, L1CAM, and ICAM-5 monoclonal antibodies, which demonstrated a band at 140kDa in fractions 8-16 (**Figure 3 c**). A similar result was obtained when polyclonal antibodies were used for ICAM-5 in fractions 8-16 at ∼130kDa. CD-63 and CD-9 were positively detected at ∼25kda in fractions 7-16 and 9-16, respectively. We were unable to detect L1CAM under reducing conditions, however we did get positive signal for L1CAM in fractions 9-16 at 220kda under non-reducing conditions. Interestingly, for ATP1A3 we observed a signal at 60kda using a monoclonal antibody in fractions 7-19. We did not detect the expected 110 kDa full-length ATP1A3 protein, in contrast to previous results using this antibody [6]. Overall, ICAM-5 was detected at the expected molecular weight, however, ATP1A3 and L1CAM, were not detected at their reported molecular weights under reducing conditions. A weak signal was detected for both L1CAM and ATP1A3 in later SEC fractions signal where signal for tetraspanins was largely absent.

**Figure 3.**
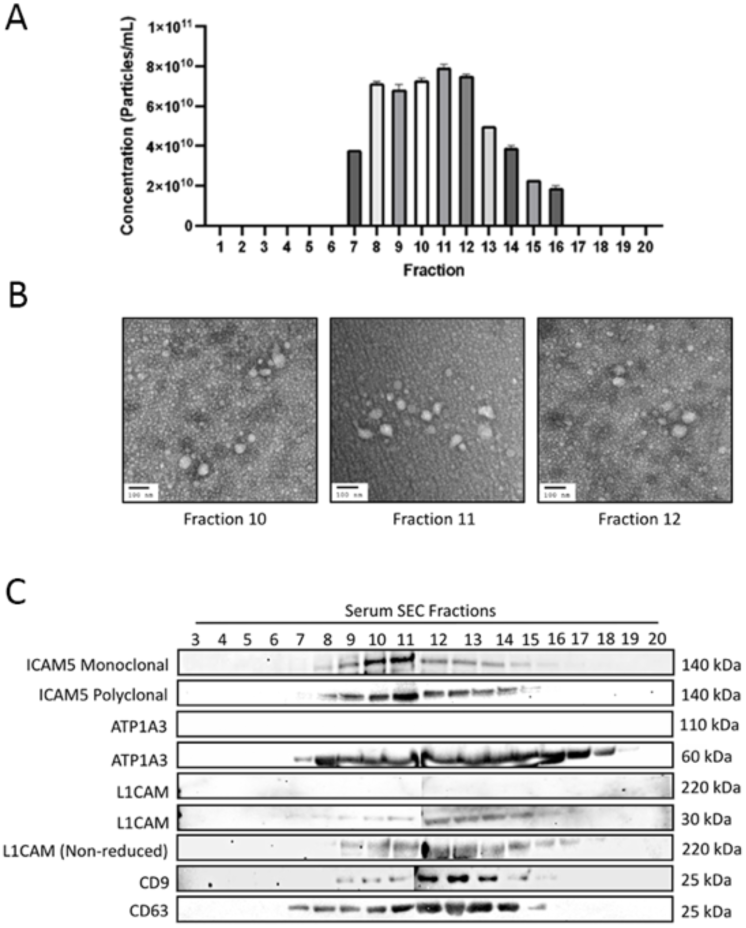
A) Nanoparticle tracking analysis (NTA) data of fractionated samples 1-20. B) Representative EM images of EVs in fractions 10, 11, and 12. C) Western blots on fractionated serum samples using SEC (fractions 3 to 20) and probed with antibodies for CD9, CD63, ICAM-5, ATP1A3, L1CAM,

Following analysis of serum, presence of ICAM-5 in EVs released in cerebrospinal fluid (CSF) was investigated. Results from NTA and EM revealed presence of EVs in only one single fraction (fraction 12) of CSF from healthy human subjects using SEC (**Figure 4 a&b**). Western blot for different SEC fractions was performed and demonstrate positive signal only in fraction 12 at 25kDa antibodies for tetraspanins (CD-9 and CD-63) (**Figure 4c**). For ICAM-5, a monoclonal antibody shows a signal at a 140kDa band in fraction 12 only. The same pattern was observed with polyclonal ICAM-5. For ATP1A3, we could not detect any band at the predicted molecular weight of 110 kDa, however similar to serum sample, a smaller molecular weight signal was detected. No signal was detected for L1CAM in EVs isolated from CSF at the predicted molecular weight.

**Figure 4.**
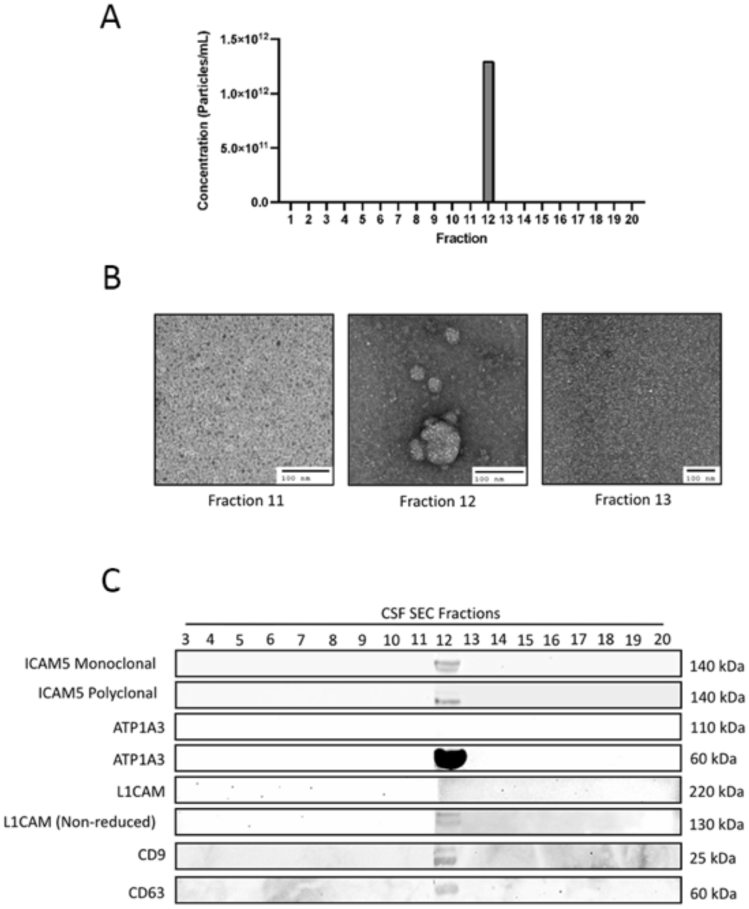
CSF EVs fractionated using size exclusion chromatography. A) NTA of 20 fractions found EVs present in only fraction 12 B) Representative EM images from fractions 11, 12, and 13. C) Western blot analysis of fractionated samples from human CSF probed with antibodies for CD9, CD63, ICAM-5, ATP1A3, and L1CAM.

### 4. Super resolution microscopy determines the quantitation of ICAM-5 positive EVs in human serum and CSF

**Direct stochastic optical reconstruction microscopy (dSTORM)** was performed to investigate whether ICAM-5 epitopes are present on the surface of EVs. This high-resolution technique enabled us to visualize and quantify specific subpopulation of EVs based on the presence a surface marker. For dSTORM analysis, EVs isolated from human serum samples were used and analyzed for the presence of ICAM-5, ATP1A3 along with total EV marker (Pan EV) (**Figure 5a**). This analysis identified EVs positive for ICAM-5 (ICAM-5+Pan-EV) and ATP1A3 (ATP1A3+Pan-EV) within the total EV population that showed a positive signal for ICAM-5 as well as ATP1A3. The ICAM-5+Pan-EV subpopulation constituted approximately 8.61% of the total EV population. In contrast, a larger proportion of EVs (21.74%) were positive for ATP1A3 (ATP1A3+Pan-EV) (**Figure 5b and d**). In EVs isolated from human CSF samples (fraction 12 of SEC), confirmed presence of both ICAM-5 and ATP1A3 on these vesicles. A higher percentage of ICAM-5+Pan-EV (20.15%) compared to the ATP1A3+Pan-EV (17.27%) was observed (**Figure 5c and d**). The results indicate there are higher number of ICAM-5 positive EVs in CSF compared to serum, while this was not observed for ATP1A3.

**Figure 5.**
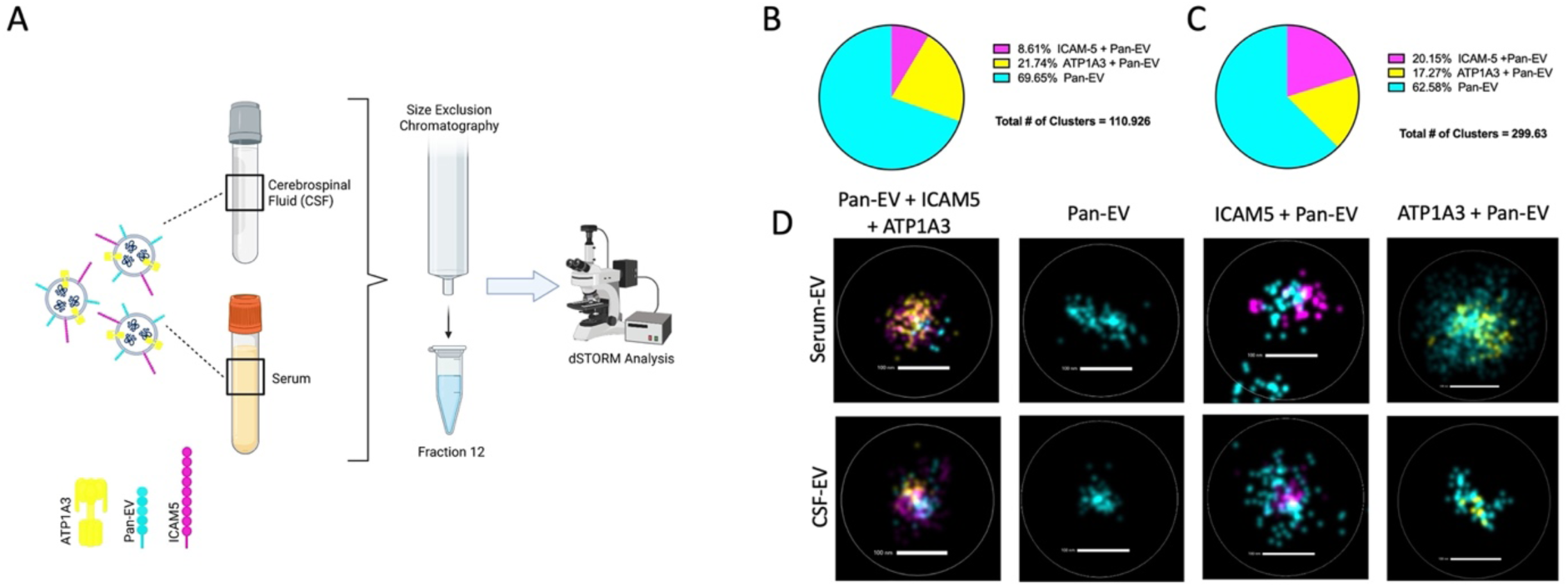
dSTORM microscopy of EVs from human serum and CSF. A) Schematic of the dSTORM experimental design (created with biorender). B) Abundance of EVs isolated from human serum. EVs positive for ICAM-5+Pan-EV markers made up 8.61%, whereas EVs positive for ATP1A3+Pan-EV made up 21.74% of the total EV clusters. C) Abundance of EVs isolated from CSF through size exclusion chromatography. D) Representative images of EVs from serum and CSF.

### 5. EVs immunoprecipitated with ICAM-5 are enriched in neuronal proteins

With confirmed presence of ICAM-5 epitopes on human serum EVs, the next objective was to determine if ICAM-5 EVs could be successfully isolated from human serum and if they contain a high concentration of neuronal proteins. To demonstrate this, total EVs from human serum samples were isolated and NDEVs were purified using biotinylated antibodies for ICAM-5, L1CAM and ATP1A3. Total EVs were included as control (**Figure 6a**). EM was performed and confirmed the presence of EVs in all groups (**Figure 6b**. Next, neuronal, glial, and EV-specific proteins were assessed in these NDEVs. Western blot was performed with antibodies for neuronal rich proteins (Tau, NSE), a glial protein (GFAP), and CD-63. The results of protein analysis in NDEVs via Western blotting demonstrate that all three markers (ICAM-5, ATP1A3, and L1-CAM) produced a positive signal for neuronal proteins. Also, ATP1A3 and L1-CAM show signal for GFAP, whereas for ICAM-5 no GFAP signal was detected. The total EVs that were loaded as controls showed the strongest signal for all the proteins (**Figure 6c**.

**Figure 6.**
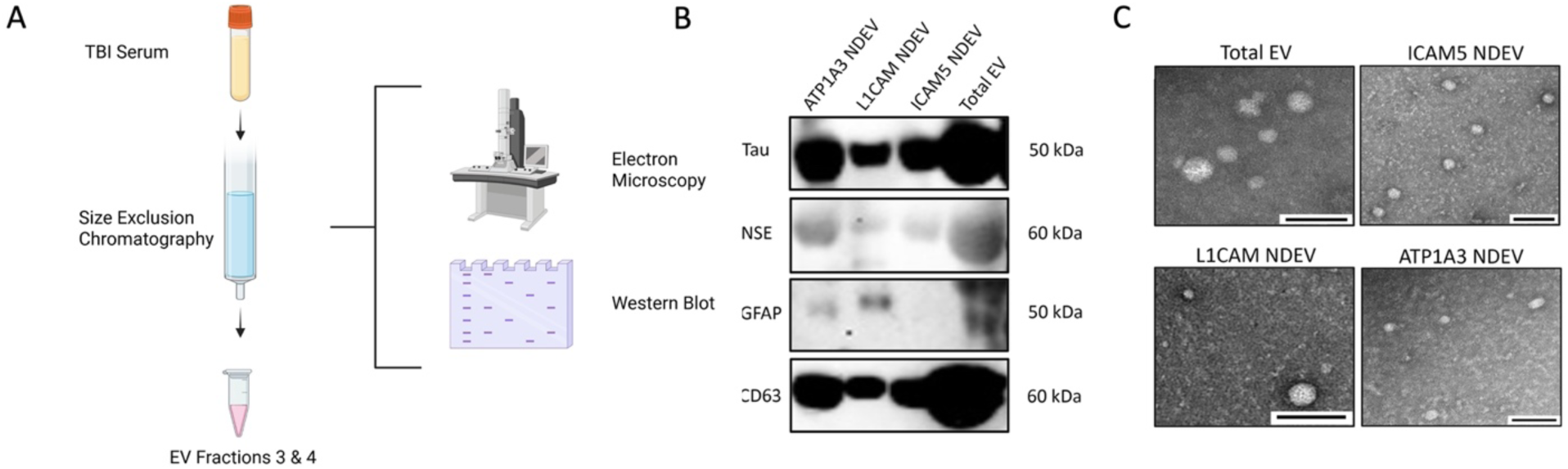
A. Schematic of the experimental design (created with Biorender). B) Western Blot of total purified EVs and NDEVs isolated using ICAM-5, L1CAM, and ATP1A3 for neuronal proteins (Tau and NSE), glial proteins (GFAP), and Tetraspanins (CD63). C) Representative images from electron microscopy (EM) of total EV samples, and EVs immunoprecipitated using ICAM-5, L1CAM, and ATP1A3 antibodies (measure bar = 100nm).

### 6. ICAM-5 derived NDEVs detects higher concentration of neuronal biomarker in serum samples from human cases of TBI

Ubiquitin C Terminal Hydrolase 1 (UCH-L1), which is an FDA approved marker for detecting TBI was assayed in ICAM-5 NDEVs and compared with total EV, L1-CAM NDEVs and ATP1A3 NDEVs. Total EVs were isolated from serum of mild TBI patients (n=10) within 24 hr of injury (**Table 1**, **Figure 7a**)). NTA was performed to confirm the relative abundance of the NDEVs between the three different antibodies which showed no significant difference in the yields of EVs between the three markers. NTA analysis also shows that NDEVs are less than 2% of the total EVs isolated from serum samples (**Figure 7b**). Quantitative ELISA shows that ICAM-5 NDEVs has the highest UCH-L1 concentration (mean = 1520 pg/mL) among the three NDEV markers (**Figure 7c**). NDEVs isolated using the ICAM-5 NDEVs had a statistically higher concentration of UCH-L1 when compared with the NDEVs isolated using L1-CAM (p = 0.0019) and ATP1A3 (p = 0.0049). Interestingly, the concentration of UCH-L1 in ICAM-5 EVs exceeded the total EV concentration of UCH-L1. The concentrations in **Figure 7** are representative of pg/mL of EV suspension and not of total serum. These results demonstrate ICAM-5 as a novel marker for enrichment of NDEVs from human serum samples that outperforms other NDEV markers in detecting neuronal injury biomarker.

**Figure 7.**
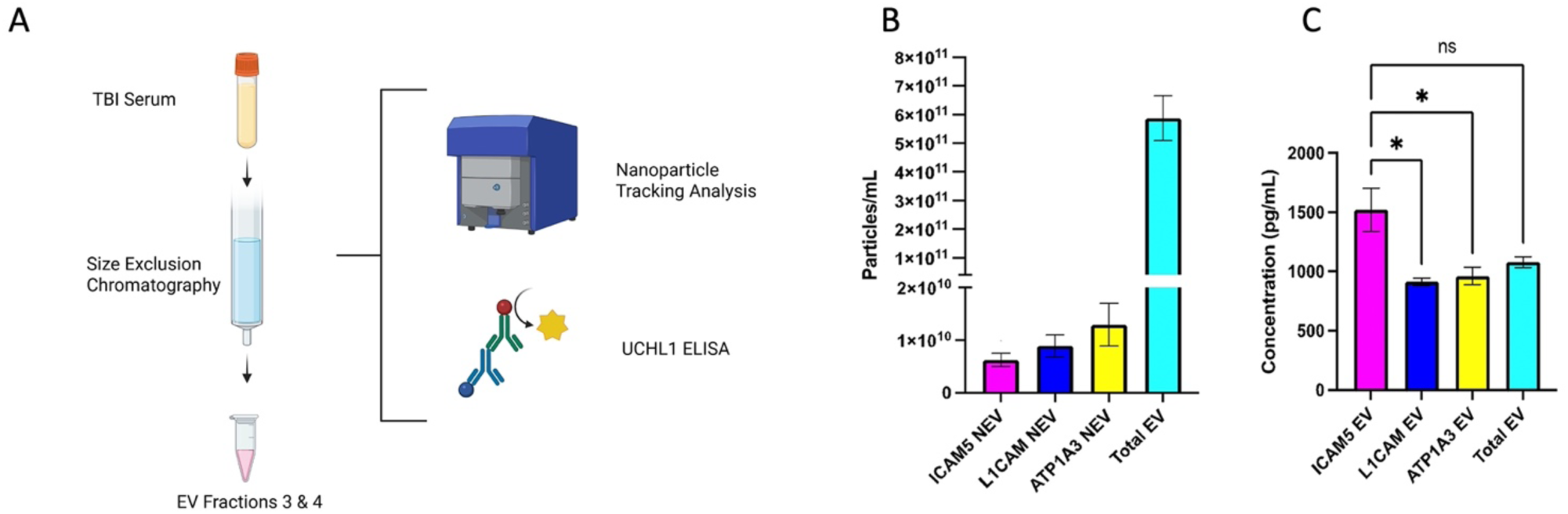
A) Schematic of experimental design for testing UCH-L1 concentration in ICAM-5 EVs isolated from patient’s serum sample with mild TBI (created with biorender). B) Nanoparticle Tracking Analysis (NTA) revealed that in comparison to total EVs, NDEVs were less than 2% of the total EVs. Comparison of UCHL1 in NDEVs, of mild TBI (mTBI) samples at 12 hours of injury, show that ICAM-5 NDEVs were able to detect the highest concentration of UCHL1 when compared to L1CAM (p = 0.0019) and ATP1A3 (p = 0.0049).

**Table 1:** Demographics of participants with a traumatic brain injury.

|  | CT Positive Scan | CT Negative Scan |
| --- | --- | --- |
| **Number of participants (N)** | 7 | 3 |
| **Sex** |  |  |
| **Male** | 6 | 2 |
| **Female** | 1 | 1 |
| **Average Age** | 49.38 | 28.67 |
| **Injury Cause** |  |  |
| **Road Traffic Accident** | 2 | 2 |
| **Incidental fall** | 4 | 0 |
| **Other non-intentional injury** | 0 | 1 |
| **Violence/assault** | 1 | 0 |

## Discussion

NDEVs are important for the identification and detection of biomarkers associated with neurodegenerative diseases and disorders, including neuronal injuries such as traumatic brain injury (TBI) and stroke. Specific markers of NDEVs are highly desired for isolation of a pure population of NDEVs from human biofluids such as serum, plasma, CSF and urine. Currently, several markers for specific isolation of NDEVs have been proposed such as L1CAM [12], ATP1A3 [4]), Neurexin (NRXN3) [13], Growth-Associated Protein (GAP) 43 and Neuroligin 3 (NLGN3)[14]. However, there is no consensus in literature for a reliable marker for NDEV isolation due to a lack of specificity of the current markers for CNS EVs and due to their presence as soluble proteins in biofluids. The primary objective of this study was to identify a novel marker that is specific to human neurons and is primarily present on the EV surface. Our study demonstrates that ICAM-5 serves as a novel and specific marker for NDEVs derived from human neurons in the CNS. It is predominantly present on the EV membrane within biofluids. ICAM-5 is also known as telencephalin, is a type-I transmembrane glycoprotein that is specifically expressed in the telencephalic neurons of the CNS [15]. ICAM-5 is the largest protein among the family of ICAM proteins with 9 extracellular domains, and is present on the dendrites and the dendritic spines of the neurons where it facilitates cellular adhesion and plays a key role in synaptic plasticity [16]. ICAM-5 can bind to β2-integrin LFA-1 on leukocytes which can result in a neuroinflammatory process in the CNS [17]. Other members of the ICAM family are expressed in many different tissues and cell types, however, expression of ICAM-5 is restricted only to neurons. This makes ICAM-5 an ideal marker for enrichment of NDEVs from biofluids.

An unbiased screening of EVs derived from the supernatant of the PNC was performed to identify potential markers for NDEVs. MS data was analyzed to identify the markers whose expression was highly restricted to the CNS. This analysis also identified ICAM-5 as a potential marker for NDEVs alongside previously reported markers for NDEV such as L1-CAM, NRXN3, and ATP1A3 and EV markers such as CD-9 and CD-81. To ensure presence of ICAM-5 on EVs derived from human biofluids, we utilized human serum samples from individuals who sustained mild TBI. We chose serum samples from mild TBI because TBI may lead to a transient leaky blood brain barriers allowing more EVs for sensitive detection [18]. The western blot results showed a specific band for ICAM-5 in the total EV samples from mild TBI patients along with presence of other neuronal markers. This confirmed that presence of ICAM-5 is in the EVs isolated from human serum samples.

For efficient enrichment of NDEVs from biofluids, the marker should be predominantly present on the NDEVs. Their presence as soluble protein or as other protein complexes will lead to poor NDEV enrichment and may lead to false positives in subsequent biomarker analysis. Recent reports suggests that L1-CAM is present in biofluids as a soluble protein and that can limit its utility as a reliable NDEV marker [6, 7]. Therefore, to confirm if ICAM-5 is an EV-associated, SEC was performed followed by western blot. SEC elutes EVs based on the size and therefore it is useful in differentiating whether the eluted protein is a soluble protein or EV associate protein. Western blots were performed for ICAM-5, L1-CAM, ATP1A3, CD-63 and CD-9 on the SEC fractions to determine the localization of these proteins with reference to the EV markers. The results showed that ICAM-5 was present from fraction 9-14 that colocalized with the tetraspanins markers suggesting that a bulk of ICAM-5 is EV associated. These results were confirmed with both a monoclonal and polyclonal antibody for ICAM-5. For ATP1A3 and L1-CAM, no signal was detected at their predicted molecular weights, however, a signal at smaller molecular weight was observed ∼50-60kDa for ATP1A3 and ∼30kDa for L1-CAM. Signal for both ATP1A3 and L1CAM was detectable in later fractions, after the tetraspanins had eluted. This may indicate that these proteins exist, at least in part, as soluble proteins. In human CSF samples, EVs were observed in only one single fraction of SEC (fraction 12) and EM confirmed this finding. This finding is notable as it contrasts with observations in other body fluids, where EVs typically elute across multiple fractions. The result, however, aligns with a previous report that described a similar phenomenon. [19]. ICAM-5 signal on western blot was detected at a molecular weight consistent with serum samples, but exclusively in fraction 12, along with the tetraspanins CD63 and CD9. However, a noticeably weaker protein signal was observed compared to serum EVs, which is expected given that CSF has a significantly lower EV concentration. [20]. For ATP1A3, a signal was detected although of a much smaller size protein compared to its predicted molecular weight in the same fraction. No signal for L1-CAM was detected in CSF EVs in any of the fractions. Presence of EVs in a single fraction may suggest presence of uniformly sized EVs in the CSF where the osmotic balance is tightly regulated. However, EM results were confirmatory that EVs were present and ICAM-5 reactivity suggest that a full length ICAM-5 is likely present in the neuronal EVs. The presence of a smaller size band for ATP1A3 in both serum and CSF fractions was observed. Several factors could explain this result; one possibility is proteolytic cleavage which is a common. Another possibility is the non-specific binding of the antibody to other isoforms or truncated versions of the protein; however, this was not investigated further in this study.

The utility of ICAM-5 as a useful marker for NDEV enrichment hinges upon its presence as a surface protein on the EVs. To evaluate this, dSTORM was performed using EV fractions from both human serum and CSF samples. We used labelled antibodies to detect ATP1A3 and ICAM-5, as well as a general marker to identify all the EVs. L1-CAM was not assayed since it was not detectable in CSF in our earlier experiments. For these studies, SEC fraction 12 was used for both serum and CSF based on our results that showed high abundance of EVs in this fraction. dSTORM experiments showed a positive detection of ICAM-5 as well as ATP1A3 for both the fractions suggesting presence of both proteins on the EV surface. Quantitative assessments were performed to calculate the relative abundance of EVs positive for ATP1A3 and ICAM-5. This analysis identified ∼ 21.74% of total EVs to be positive for ATP1A3 whereas ∼ 8.61% EVs were positive for ICAM-5 in serum samples. In CSF samples, ATP1A3 positive EVs reduced to ∼ 17% whereas ICAM-5 positive EVs increased to ∼20.15%. The results for ICAM-5 followed the predicted pattern, with a lower proportion of NDEVs relative to total EVs in serum versus a higher proportion in CSF. In contrast, ATP1A3 exhibited an unexpected trend, showing a higher number of ATP1A3 EVs in serum than in CSF.

After confirming that ICAM-5 is present on EV surface, we wanted to confirm whether ICAM-5 can be used to isolate NDEVs from human serum samples and whether they are rich in proteins of neuronal origin. To evaluate this, we used control human serum samples and performed immunoprecipitation with ICAM-5, ATP1A3 and L1CAM antibody. The western blot results showed a significant enrichment of neuronal proteins NSE and Tau in NDEVs enriched with ICAM-5. The EVs eluted with ATP1A3 and L1-CAM also showed enrichment of neuronal protein. GFAP which is a specific marker of astrocytes was not detected in ICAM-5 enriched EVs however GFAP was detected in both ATP1A3 and L1CAM eluted EVs. This indicates that ICAM-5 is a specific marker for neurons. Finally, to validate the clinical utility of ICAM-5 derived NDEVs, we evaluated the expression of an FDA approved biomarker for TBI neuronal marker UCH-L1 in a cohort of mild TBI patients. The samples for this cohort were collected within the 12 hr post injury. UCH-L1 is a neuronal specific marker and is known to be elevated after TBI within 12 hrs post injury [21]. We compared the expression of UCH-L1 in NDEVs isolated with ICAM-5, ATP1A3 and L1CAM. Our results showed a significantly higher expression of UCH-L1 in NDEVs eluted using ICAM-5 antibodies compared to ATP1A3 and L1CAM. NTA analysis showed no significant differences in the numbers of NDEVs from either of the three markers. This suggest that ICAM-5 EVs were likely of a pure neuronal EVs representing higher concentration of neuronal proteins compared to ATP1A3 and L1CAM. the percentage of EVs positive for ICAM-5 observed after immunoprecipitation was found to be much less compared to the data from dSTORM analysis. This may indicate a potential for further optimization of the immunoprecipitation protocol for better recovery.

## Conclusions

This study identified ICAM-5 as a novel specific marker for purification and enrichment of NDEVs from human biofluids. To our knowledge this is the first study to compare the known markers of NDEV isolation demonstrating a direct comparison of these markers for their specificity to CNS and their potential in isolation of neuronal EVs. Our results suggest ICAM-5 has a potential to be clinically relevant markers for discovery and detection of neuronal biomarkers. Further studies are needed to evaluate its potential for its widespread adoption in research as well as clinical setting.

### Limitations

This study has a few limitations. First, while we used equal amounts of ICAM-5, L1CAM and ATP1A3 antibodies for all immunoprecipitation assays, we did not perform an optimization of antibody concentration and incubation times. Future studies could explore these variables to potentially improve the efficiency of NDEV isolation. Second, the sample size for testing UCH-L1 in clinical TBI samples was relatively small. Although our results show a promising trend, these findings should be replicated in a larger, more diverse cohort to confirm the reproducibility and broader applicability of our data. Finally, our quantitative analysis focused exclusively on UCH-L1 as a single biomarker. Future research should include a broader range of biomarkers across various neurological conditions. This would provide further evidence to support the utility of ICAM-5 as a robust marker for NDEVs from biofluids.

## Methods

### Cell Culture

Human primary neuronal cells (PNC), media and growth factors for optimized cell growth were acquired from Neuromics Inc and cultured as per recommended protocol. Immunofluorescence assays performed using neuronal specific (neuronal specific enolase, NSE, Sigma Aldrich, MAB324) and glial cell specific (glial fibrillary acidic protein, GFAP, Santa Cruz, sc-71143). Dependent on experimental conditions and 24-hr prior to any NDEV isolation, serum-rich PNC culture media was replaced with serum-free media to remove EVs associated with bovine serum. For MS analysis, total EVs from 50 ml of serum free cell culture media were isolated using ExoQuick-TC (System Biosciences Inc).

### Mass Spectrometry

Samples were analyzed by LC-MS using Nano LC-MS/MS (Dionex Ultimate 3000 RLSCnano System, Thermofisher) interfaced with Eclipse (Thermofisher). Raw data were analyzed with predicted libraries from specified databases and for library-free search using DIA NN 1.8.1 with recommended settings. MS analysis was performed by System Biosciences Inc, CA. MS for the EVs isolated from supernatants of neuronal cells yielded more than 3,000 protein hits which was compared to the list of proteins in human protein atlas (proteinatlas.org) under the heading “elevated proteins in the human brain.” These 63 proteins were then analyzed for transmembrane glycoproteins from literature reports.

### Samples

Human serum samples (1mL) were obtained through a commercial vendor (BIOIVT LLC, HUMANSRM-0101). Mild TBI serum samples were obtained through the clinical biomarkers core at the Military Traumatic Brain Injury Initiative (MTBI^2^) biorepository, Uniformed Services University of Health Sciences (USUHS).

### EV isolation methods

#### EV isolation from TBI samples

IZON qEV original Gen 2 70nm (IZON Science, CAT#: ICO-70) columns for SEC were used to isolate total EVs from 250uL of 3 vials of pooled mTBI serum (Figure 2). The sample was centrifuged at 1500xg for 10min to remove cellular debris before EV isolation. 2.5mL of 1XPBS was used to flush the column before the addition of serum. The serum was let to fully saturate into the resin before adding 2.25 mL of 1XPBS and the buffer was let to run through. 400 µL of 1XPBS was then added to the column and the EV flow-through was collected. 400 µuL of 1XPBS was added 5 times to create 5 EV fractions. Fractions 3 and 4 were pooled to create a total volume of 800uL before immunoprecipitation with ICAM-5 biotinylated. 200 µL of total EVs from fractions 3 and 4, were used for immunoprecipitation. Immunoprecipitation from total EVs yielded a total of 200 µL ICAM-5 immunoprecipitated EVs which were then analyzed for the presence of EV-specific markers using the Exo-Check Exosome Antibody Neruo-Array Standard (Systems Biosciences, CAT#: EXORAY500a-8), according to the manufacturer’s recommended protocol, or analyzed on Western Blots probed with rabbit-anti-ICAM-5 polyclonal (Boster Biological Technologies, CAT#: A07187-2) to analyze the presence of ICAM-5 in the samples.

#### EV isolation for characterization Assay using Size Exclusion Chromatography

200μl of human serum or CSF (BioIVT) were used for isolation via size exclusion chromatography (SEC) with IZON qEV single use SEC columns (IZON, CAT#: ICS-70) (**Figure 3**, **figure 4 and figure 5**). Samples were first centrifuged at 10,000 x g for 10 minutes to remove any large particulates and cells. The supernatant was then transferred to a new microcentrifuge tube). The qEV column and sample buffer were equilibrated to room temperature (18°C-24°C). The top cap of the column was removed, followed by the removal of the bottom cap. A buffer volume collection vessel was placed below the column. The column was flushed with 3 ml of 1xPBS two times according to the manufacturer’s recommended protocol. 150µl of sample was loaded into the loading frit and was allowed time to fully saturate into the resin. Then, 720µl of 1xPBS was loaded into the column, to equal a total of 870µl of fluid. A microcentrifuge tube was placed below the column, and a background fraction was collected. After the background was collected, a new collection vessel was placed below the column. The flow through was collected 20 times for 20 separate fractions by adding 170µl of 1xPBS to the column for each fraction. For each fraction, the flow-through had to completely stop before adding the next 170µl of 1xPBS. This process was repeated for 20 fractions. EV fractions 1-20 were then analyzed using Nanoparticle tracking analysis (NTA), electron microscopy (EM) to quantify and determine the presence of EVs within each fraction. Samples 3-20 were used for Western Blots to characterize ICAM-5. The following primary antibodies were used for western blotting for the characterization assay: rabbit-anti-ICAM-5 monoclonal (abcam, CAT#: ab302899), rabbit-anti-ICAM-5 polyclonal (Boster Biological Technologies, CAT#: A07187-2), mouse-anti-CD-9 monoclonal (Invitrogen, CAT#: 10626D), mouse-anti-C63 monoclonal (Thermo Fisher, CAT#: 10628D), mouse-anti-ATP1A3 monoclonal (Invitrogen, CAT#: MA3-915), and mouse-anti-CD171 monoclonal (Invitrogen, CAT#: 13-1719-82). CSF fraction 12 and serum fraction 12 were used for subsequent dSTORM analysis.

#### EV isolation for neuronal protein enrichment validation

Total EVs from 200 µL of human serum (BioIVT) was isolation of using the Plasma/Serum Exosome Purification and RNA Isolation Kits (Norgen Biotek, CAT #: 58300) according to the manufacturer’s recommended protocol (**Figure 6**). Nuclease-free water was added to each serum sample to bring the volume of the samples up to 1mL. From this protocol, 600μl of purified exosomes was obtained. 200μl of purified EVs were then used for immunoprecipitation with ICAM-5, L1CAM, and ATP1A3, subsequent testing for the presence of EVs with Electron Microscopy and western blotting for the presence of neuronal and glial proteins. The following primary antibodies were used for western blotting: rabbit-anti-Tau polyclonal (Santa Cruz, CAT#: SC-5587), mouse-anti-NSE monoclonal (Sigma Aldrich, CAT#: MAB324), mouse-anti-GFAP monoclonal (Santa Cruz, CAT#: SC-71143), and mouse-anti-C63 monoclonal (Thermo Fisher, CAT#: 10628D).

#### EV isolation for UCH-L1 ELISA assay

IZON qEV original Gen 2 70nm (IZON Science, CAT#: ICO-70) columns for SEC were used to isolate total EVs from 200uL of mTBI serum. Samples were centrifuged at 1500xg for 10min to remove cellular debris before EV isolation. 2.5mL of 1XPBS was used to flush the columns before the addition of 200 µL of serum. The serum was let to fully saturate into the resin before adding 2.25 mL of 1XPBS and the buffer was let to run through. 400 µL of 1XPBS was then added to the column and the EV flow-through was collected. 400 µL of 1XPBS was added 5 times to create 5 EV fractions. Fractions 3 and 4 were pooled to create a total volume of 800 µL, which was then divided into 4 treatment groups: total EV, ICAM-5 immunoprecipitated NDEV, L1CAM immunoprecipitated NDEV, and ATP1A3 immunoprecipitated NDEV. Multiplex commercial kits (Thermo Fisher Sci., CAT# EPX010-10420-901) were used to evaluate the protein concentration of GFAP (Thermo Fisher Sci., CAT# EPX010-12336-901) and UCHL1 (Thermo Fisher Sci., EPX010-12338-901) in purified total EV and samples immunocaptured with ICAM-5, L1CAM, and ATP1A3, according to the manufacturer’s recommended protocol. All samples were duplicated.

#### Immunoprecipitation

200 μl of total purified EV were combined with 4 μg of biotinylated antibody for ICAM-5 (Boster biologics, CAT#: A07187-2), L1CAM (Invitrogen, CAT#: 13-1719-82), or ATP1A3 (LSBio, CAT#: LS-C500359-50), in a total of 50 μl of 3% BSA per tube. The suspensions were incubated overnight at 4°C with constant mixing. 15 μl of streptavidin-agarose Ultralink resin (Thermo Fisher Scientific, CAT#: 53114) in was added to 25 μl of 3% BSA, which was then added to the EV suspension and incubated for 1 hour at 4°C with continuous mixing. After incubation, samples were centrifuged at 200 × g for 10 minutes at 4°C. The supernatant was removed and the pellet was resuspended in 200 μl of 0.1 M glycine-HCl (pH 2.5) by mixing for 10-15 seconds, and then centrifuged at 4,500 × g for 10 minutes at 4°C to detach the targeted EVs from the bead-antibody complex. The supernatants were then transferred to clean tubes containing 25 μl of 3% BSA and 15 μl of 1M TRIS-HCl.

#### Nanoparticle Tracking Analysis (NTA)

Nano-particle tracking analysis was performed using the Nanoparticle Tracking Analyzer ZetaView TWIN (ZetaView). Total EV fractions isolated using SEC and immunoprecipitated NDEV samples were diluted at 1/5000 and 1/500, respectively. All dilutions were made in 5 mL of UltraPure Distilled Water. 2 mL of the diluted samples were loaded with a syringe into the analyzer. The Brownian motion of EV particles was analysed under a laser wavelength of 488 nm for 1 cycle in 11 positions. ZetaView 8.06.01 was used to analyse the recorded videos with the following settings: sensitivity 80, shutter 100, and frame rate 30. Each sample recorded was performed in duplicates.

#### Electron Microscopy (EM)

EVs were visualized using Electron Microscopy at the Biomedical Instrumentation Center (BIC) at the Uniformed Services University using a Transition Electron Microscope (JEOL). EVs were placed on copper carbon coated grids (Electron Microscopy Sciences, CAT#: FCF400-Cu-50) for 5 min before negative staining with 3% uranyl acetate. Before the addition of EVs, copper grids were made hydrophilic by a 3-minute exposure to a glow discharge.

#### Western Blotting

16µl of sample from CSF and serum were lysed using 10 µl of RIPA buffer (G Biosciences, CAT#: 786-489) to prepare for Western Blotting. The lysed sample was then combined with 4 µl of NuPage sample reducing agent (Invitrogen, CAT#: NP0009) and 10 µl LDS Sample Buffer (Thermo Fisher Scientific Inc., CAT#: 84788), and loaded on NuPage 4-12% Bis-Tris Gels (Invitrogen, CAT#: NP0321BOX). The gels were then wet transferred onto 0.45-μm nitrocellulose membranes (Bio-Rad CAT#: 1620145), and blocked using 5% nonfat dry milk in 1x TBS-Tween20 for 1 hour at room temperature before being incubated with primary antibodies overnight at 4°C with constant shaking. The membranes were washed with 1x TBS-T 3 times for 5 min each at 100 rpm and further incubated for 1 hour in 5% non-fat dry milk with the following appropriate secondary antibodies: Goat-anti-rabbit IgG (Abcam, CAT#: AB7090) and Goat-anti-mouse IgG (Abcam, CAT#: AB47827). Immunoreactivity was captured using Western Bright Sirius substrate (Advansta, CAT#: R-03027-D1) and all images were captured using the iBright FL1000.

#### dSTORM Analysis

To confirm the localization of markers on the EV surface, direct stochastic optical reconstruction microscopy (dSTORM) was conducted by the Extracellular particle Characterization and Enrichment Lab (EXCEL) at Johns Hopkins University. Single molecule localization microscopy was performed using the Nanoimager (Oxford Nanoimaging) with the EV profiler 2 kit (Oxford Nanoimaging) with Tetraspanin Trio capture (Oxford Nanoimaging) for CSF samples and CD81 capture for plasma samples, following the manufacturer’s instructions. Briefly, 9 μL samples were incubated on a chip for 60 minutes, washed, and labeled with fluorescent antibodies for ICAM-5-647, ATP1A3-555, and Pan EV-488 staining (Oxford Nanoimagin) for 60 minutes. Samples were imaged using the AutoEV mode with Pan-EV +Teraspanins settings, and AI EV profiling data analysis tool. Conjugated antibodies (ICAM-5 monoclonal antibodies (R&D Systems, CAT#: MAB1950) were labeled using the Alexa Fluor 647 Conjugation Kit (Abcam, CAT#: 269823) as per the manufacturer’s recommended protocol. ATP1A3 monoclonal antibodies (Invitrogen, CAT#: MA3-915) Alexa Fluor 555 Conjugation Kit (Abcam, CAT#: 269820) as per the manufacturer’s recommended protocol. Microscopy was performed on EV fraction 12 in serum samples and Fractions 12 in CSF samples.

#### Statistical Analysis

Statistical analysis was performed using Graphpad Prism 10. Demographic characteristics from mTBI subjects were compared between CT positive and CT negative groups using independent 2-tailed T-tests, and Chi-squared tests. Protein concentrations obtained through ELISA were analyzed using one-way ANOVA. Results from NTA were analyzed using ANOVA. In all analyses, p ≤ 0.05 was considered significant.

## Institutional Review Board Statement

The De-identified mild TBI samples were requested from the Center for Neuroscience and Regenerative Medicine’s biorepository, USUHS. The controls serum samples were commercially acquired. The Institutional review board of Uniformed Services University, reviewed the use of de-identified samples and approved these samples to be used in this study.

## Informed Consent Statement

Informed consent was obtained from all subjects whose blood samples were used in this study.

## Data Availability Statement

All the data associated with this study is presented in the main manuscript and the supplementary data section.

## Disclaimer

The opinions and assertions expressed herein are those of the authors and do not necessarily reflect the official policy or position of the Uniformed Services University or the Department of War or the U.S. Government. The opinions and assertions expressed herein are those of the authors and do not necessarily reflect the official policy or position of the Henry M. Jackson Foundation for the Advancement of Military Medicine. The authors thank all participants for their time and contributions to this research.

