## Supplementary table 2 for "ICAM-5: A Novel Marker for Neuronally Derived Extracellular Vesicles"

**Supplementary table 2**: List of proteins identified after analysis with proteins enriched in brain using proteinatlas.org.

| **ACTL6B** | **COL20A1** | **FSTL5** | **LINGO2** | **SERPINI2** |
| --- | --- | --- | --- | --- |
| **ALK** | **COL26A1** | **GPC2** | **MAB21L1** | **SEZ6L** |
| **ARMC3** | **COL2A1** | **GPRIN1** | **NCAN** | **SLIT1** |
| **ATCAY** | **DBH** | **H2BC26** | **NELL1** | **ST8SIA2** |
| **BHMT** | **DCC** | **H4C1** | **NKAIN1** | **ST8SIA3** |
| **C10orf90** | **DCX** | **HPDL** | **OLFM3** | **SYT5** |
| **C1QL4** | **DPYSL5** | **ICAM5** | **PPM1E** | **TFAP2B** |
| **CAMKV** | **ELAVL2** | **IGFALS** | **PZP** | **TH** |
| **CBLN4** | **ELAVL4** | **IGFBPL1** | **RAB39A** | **TMEFF1** |
| **CELSR3** | **FAT3** | **INA** | **RAB3C** | **TNR** |
| **CNTN2** | **FBLL1** | **KRT31** | **RELN** | **VGF** |
| **CNTNAP2** | **FGL1** | **LAMA1** | **RTL1** | **XKR7** |
| **CNTNAP4** | **FSD1** | **LCN1** | **SCG3** | **ZNF469** |
